# Limited uptake of essential amino acid is effective for cancer therapy in metabolic networks of integrated constraint-based models

**DOI:** 10.1101/281253

**Authors:** Takeyuki Tamura

## Abstract

Nam models are context-specific genome-scale metabolic models for nine types of cancer and corresponding normal cells reconstructed from RECON 2, genetic mutation information, and expression profile data. In this study, computational experiments were conducted using Nam models to find effective ratios of uptake reactions such that cancer cells do not grow while normal cells grow. The original Nam models were extended to consider interactions between cancer and normal cells using an approach developed for representing host-microbe interactions. When uptake ratio of only single reaction was allowed to change at a time, the results of computational experiments showed that every essential amino acid had effective ranges for almost all nine types of cancer, while the other uptake reactions rarely had such effective ranges.

## 1 Background

In the 1920s, Otto Warburg et al. showed that cancer cells metabolize approximately ten times more glucose to lactate than normal cells in aerobic conditions [6]. This increase in aerobic glycolysis in cancer cells has often been misunderstood as adaptation to dysfunctional mitochondria. However, it was later understood that mitochondrial function is not impaired in most cancer cells [14]. On the contrary, recent studies have shown that altered metabolism can trigger cancer via oncometabolites [16, 17]. For example, isocitrate dehydrogenase (IDH) mutations lead to a gain-of-function by enabling a new metabolic enzymatic reaction that produces 2-hydroxyglutarate (2-HG) from *α*-ketoglutarate. High concentration of this oncometabolite 2-HG inhibits binding of *α*-ketoglutarate (*α*-KG) to histone demethylase, thereby blocking cell differentiation [7, 16]. On the other hand, mutations on succinate dehydrogenase (SDH) and fumarate hydratase (FH) lead to loss-of-function of reactions that consume succinate and fumarate, respectively. The accumulated succinate and fumarate act as competitive inhibitors of *α*-KG-dependent oxygenases that regulate the hypoxia-inducible factor (HIF) oncogenic pathway [5, 10, 15, 17].

The aim of this study is analyze, computationally, strategies to control cancer cells utilizing metabolic differences between cancer and normal cells. Flux Balance Analysis (FBA) is one of the most widely used mathematical models to simulate metabolism. In conventional FBA, a pseudo-steady-state is assumed where the sum of incoming flows is equal to the sum of outgoing flows for each internal metabolite. Based on FBA and Boolean representations between genes and reactions, genome-scale constraint-based models have been constructed. iAF1260 and iJO1366 are genome-scale reconstructions of the metabolic network in Escherichia coli K-12 MG1655 for 1260 and 1366 genes, respectively [2, 9].

RECON 1 and RECON 2 are comprehensive and high-quality knowledge-based reconstructions of human metabolism for 1496 and 1789 genes, respectively [1, 13]. Based on RECON2 and genetic mutation information collected from more than 1700 cancer genomes, Nam et al. reconstructed context-specific genome-scale metabolic models, of nine types of cancer cells and the corresponding normal cells that are represented in the constraint-based model [8].

In this paper, we study strategies to control cancer cells using Nam models by adding nutrient uptake constraints that lead to cancer cells not growing while normal cells grow. In contrast with chemotherapy, we focus on metabolites that are uptaken in the original Nam model. There are 85 uptake exchange reactions in Nam models. Since metabolites secreted by cancer (or normal) cells may affect normal (or resp. cancer) cells in actual therapy, it may be necessary to develop an integrated model that can simulate cancer and normal cells at the same time. We developed an integrated model of cancer and normal cells using the framework developed in [12] for modeling host-microbe interactions.

## 2 Results

Computational experiments showed that, for each of the nine types of cancer in Nam models, at least eight of the nine essential amino acids (EAA) had effective ranges of uptake ratios for both independent models and integrated models. In total, the limited uptake of EAA was effective for 93.8% of the cases for both models. The other uptake reactions had effective ranges for 12.6% of the cases in the independent model, and 5.6% of the cases in the integrated model (See Table 1). All controllable cases are listed in Tables A1 and A2 for integrated and independent Nam models, respectively, where “1‘ indicates “controllable” and “0‘ indicates “not controllable”.

**Table 1.** Cancer-specific controllability by essential amino acids (EAA) and other uptake reactions for the independent and integrated Nam models.

| (A) Independent model |  |  | (B) Integrated model |  |  |
| --- | --- | --- | --- | --- | --- |
| Cancer type | EAA | Others | Cancer type | EAA | Others |
| Lung ADC | 9/9 ( 100%) | 12/76 (15.8%) | Lung ADC | 9/9 ( 100%) | 5/76 (6.6%) |
| Breast | 8/9 (88.9%) | 6/76 ( 7.9 %) | Breast | 8/9 (88.9%) | 2/76 (2.6 %) |
| Gastric | 9/9 ( 100%) | 6/76 ( 7.9 %) | Gastric | 9/9 ( 100%) | 4/76 (5.3 %) |
| Kidney | 8/9 (88.9%) | 5/76 ( 6.6 %) | Kidney | 8/9 (88.9%) | 1/76 (1.3 %) |
| Leukemia | 9/9 (88.9%) | 18/76 (23.7 %) | Leukemia | 9/9 (88.9%) | 8/76 (10.5 %) |
| Liver | 8/9 (99.9%) | 9/76 (11.8 %) | Liver | 8/9 (88.9%) | 7/76 (9.2 %) |
| Lung SCC | 8/9 (88.9%) | 5/76 ( 6.6 %) | Lung SCC | 8/9 (88.9%) | 1/76 (1.3 %) |
| Ovarian | 9/9 ( 100%) | 17/76 (19.8 %) | Ovarian | 9/9 ( 100%) | 7/76 (9.2 %) |
| Pancreas | 8/9 (88.9%) | 8/76 (10.5 %) | Pancreas | 8/9 (88.9%) | 3/76 (3.9 %) |

Since Table 1 shows that controlling integrated Nam models is more difficult than independent Nam models, we focus on integrated Nam models hereafter. Fig. 1 shows the effective ranges of the nine EAA for the nine types of cancer in integrated Nam models. Fig. 1 (A) shows that L-histidine was effective for each of the nine types of cancer when the uptake ratio was limited to 10 to 20 % of the maximum uptake ratio. Similarly, Fig. 1 (B)(C)(E)(G)(I) show that L-isoleucine, L-leucine, L-methionine, L-threonine, L-valine were effective for each of the nine cancers when the uptake ratio was limited to approximately 20-40%, 40-85%, 10-20%, 20-50%, and 20-50% of the maximum uptake ratio, respectively. Fig. 1 (D) shows that the L-lysine was ineffective for breast, kidney, liver, and lung SCC cancers. However, it was effective for lung ADC and gastric cancers when the uptake ratio was limited to 40-90%. It was also effective for leukemia, ovarian and pancreas cancers, when the uptake ratio was limited to 5-40%. Fig. 1 (F) shows that L-phenylalanine was effective for lung ADC, breast, gastric, kidney, leukemia, and ovarian cancers when the uptake ratio was limited to approximately 20-40%. It was also effective for liver and lung SCC cancers when the uptake ratio was limited to approximately 5-20% of the maximum uptake ratio. However, it was not effective for pancreas cancer. Fig. 1 (H) shows that L-tryptophan was effective for each of the nine cancers, but its effective ranges were very small. It was effective only when the uptake ratio was limited to approximately 2-3% of the maximum uptake ratio.

**Figure 1:**
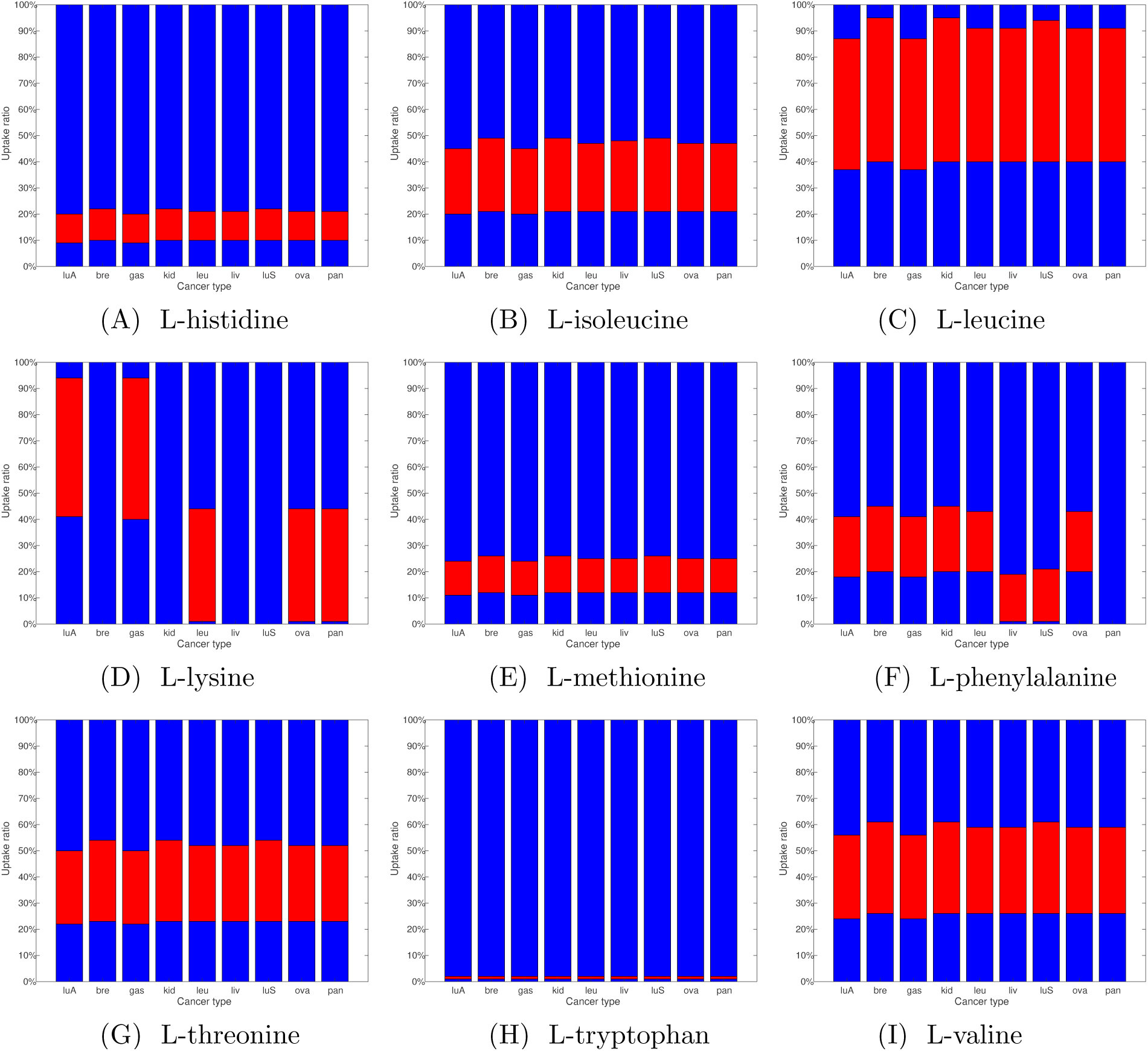
Red bars represent effective ranges of uptake ratio of each EAA for the nine types of cancer for the integrated NAM models. luA:lung adenocarcinoma(ADC), bre:breast, gas:gastric, kid: kidney, leu: leukemia, liv:liver, luS:lung squamous(SCC), ova: ovarian, pan: pancreas.

## 3 Discussion

In this section, we discuss the relationship between the results of computational experiments and the network topology of Nam models. Fig. 2 shows a toy example of a topology around an uptake exchange reaction of Nam models. Rectangles are reactions and circles are compounds. EX Rc(*i*) and EX Rn(*i*) represent the *i*th exchange uptake reaction of cancer and normal cells, respectively. In Nam models, every exchange uptake reaction has only one neighboring compound, which is represented as EX Cc(*i*) for cancer cells and EX Cn(*i*) for normal cells in Fig. 2. Although the degrees of EX Rc(*i*) and EX Rn(*i*) are one for all *i*, the degrees of EX Cc(*i*) and EX Cn(*i*) are more than one for most *i*. For any *i*, either EX Cc(*i*) or EX Cn(*i*) is not adjacent to the reaction of biomass objective function in Nam models.

**Figure 2.**
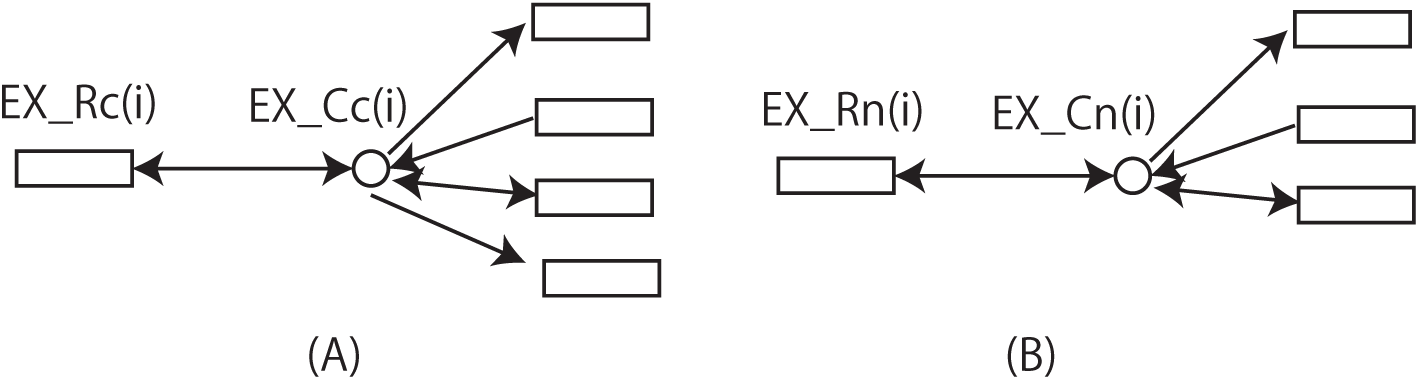
Toy examples of topology around exchange uptake reactions in Nam models. (A) *d*(*EX Cc*(*i*)) = 5, (B) *d*(*EX Cn*(*i*)) = 4.

Let *d*(*v*) represent the degree of *v*. Since *d*(*EX Rc*(*i*)) and *d*(*EX Rn*(*i*)) are one for any *i*, we focus on *EX Cc*(*i*) and *EX Cn*(*i*). We analyze the relationship between *{EX Cc*(*i*), *EX Cn*(*i*)*}* and the controllability of the exchange uptake reactions by comparing the following three algorithms.

### AlgoDiff

1. Sort exchange uptake reactions by diff= *|d*(*EX C*_*c*_(*i*)) *- d*(*EX C*_*n*_(*i*))|;
2. if diff are the same, sort by deg c=*d*(*EX C*_*c*_(*i*)).
3. if deg c are the same, sort by deg n= *d*(*EX C*_*n*_(*i*)).
4. Choose *k*-top exchange uptake reactions as “controllable”.

### AlgoDeg

1. Sort exchange uptake reactions by deg c=*d*(*EX C*_*c*_(*i*)).
2. if deg c are the same, sort by deg n= *d*(*EX C*_*n*_(*i*)).
3. Choose *k*-top exchange uptake reactions as “controllable”.

### AlgoEAA

1. Choose EAAs as “controllable”.

Since there are nine EAAs, we set *k* = 9 to compare AlgoEAA with AlgoDiff and AlgoDeg. Table 2 shows the prediction accuracy for “controllability” of AlgoDiff, AlgoDeg and AlgoEAA for the nine types of cancer of Nam models. In total, the prediction accuracies for the independent models were 42.0%, 58.0%, and 93.8% for AlgoDiff, AlgoDeg, and AlgoEAA, respectively. Similarly, the prediction accuracies for the integrated models were 37.0%, 56.8%, and 93.8% for AlgoDiff, AlgoDeg, and AlgoEAA, respectively.

**Table 2.** Prediction accuracy for controllability by AlgoDiff, AlgoDeg, and AlgoEAA for the independent model and integrated model.

(A)Independent Model
|  | AlgoDiff | AlgoDeg | AlgoEAA |
| --- | --- | --- | --- |
| Lung ADC | 7/9 (77.8%) | 4/9 (44.4%) | 9/9 ( 100%) |
| Breast | 3/9 (33.3%) | 5/9 (55.6%) | 8/9 (88.9%) |
| Gastric | 6/9 (66.7%) | 6/9 (66.7%) | 9/9 ( 100%) |
| Kidney | 4/9 (44.4%) | 5/9 (55.6%) | 8/9 (88.9%) |
| Leukemia | 3/9 (33.3%) | 8/9 (88.9%) | 9/9 ( 100%) |
| Liver | 5/9 (55.6%) | 5/9 (55.6%) | 8/9 (88.9%) |
| Lung SCC | 3/9 (33.3%) | 5/9 (55.6%) | 8/9 (88.9%) |
| Ovarian | 2/9 (22.2%) | 5/9 (55.6%) | 9/9 ( 100%) |
| Pancreas | 1/9 (11.1%) | 4/9 (44.4%) | 8/9 (88.9%) |
| total | 34/81(42.0%) | 47/81(58.0%) | 76/81(93.8%) |

(B)Integrated Model
|  | AlgoDiff | AlgoDeg | AlgoEAA |
| --- | --- | --- | --- |
| Lung ADC | 7/9 (77.8%) | 3/9 (33.3%) | 9/9 ( 100%) |
| Breast | 2/9 (22.2%) | 5/9 (55.6%) | 8/9 (88.9%) |
| Gastric | 6/9 (66.7%) | 6/9 (66.7%) | 9/9 ( 100%) |
| Kidney | 3/9 (33.3%) | 5/9 (55.6%) | 8/9 (88.9%) |
| Leukemia | 2/9 (22.2%) | 8/9 (88.9%) | 9/9 ( 100%) |
| liver | 5/9 (55.6%) | 5/9 (55.6%) | 8/9 (88.9%) |
| Lung SCC | 2/9 (22.2%) | 5/9 (55.6%) | 8/9 (88.9%) |
| Ovarian | 2/9 (22.2%) | 5/9 (55.6%) | 9/9 ( 100%) |
| Pancreas | 1/9 (11.1%) | 4/9 (44.4%) | 8/9 (88.9%) |
| total | 30/81(37.0%) | 46/81(56.8%) | 76/81(93.8)% |

The best prediction accuracy of AlgoDiff is 77.8% for Lung ADC for both independent and integrated models. Similarly, the best prediction accuracy of AlgoDeg is 88.9% for Leukemia for both independent and integrated models. It is observed that, in some cases, topological parameters such as diff and deg c can be highly correlated with “controllability”. However, the prediction accuracy of AlgoEAA is much higher than that of AlgoDiff and AlgoDeg for both independent and integrated models.

## 4 Conclusions

Our computational experiments using Nam models and FBA showed that change in the uptake ratio of an essential amino acid can result in a low growth rate of cancer cells and a normal growth rate of normal cells for 93.8% of the cases for both integrated and independent models. The control effect of EAAs in the experiments was remarkable, since other exchange uptake reactions only had such effective range of uptake ratio for 12.6% and 5.6% of the cases for independent and integrated models, respectively. Although the findings of this study are based only on computational experiments, they may help to develop a new paradigm for cancer therapy.

## 5 Methods

To computationally predict cell growth using FBA, the biomass objective function needs to be determined. This function describes the rate at which all biomass precursors are made in the correct proportions [3]. As cancer cells are known to maximize their proliferation rate [4, 11], the biomass formation is maximized in cancer cells in Nam models. However, normal cells may have different objective functions depending on growth signals [14]. Normal cells in Nam models maximize

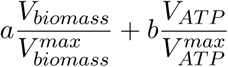

where *a* = *b* = 1 were used in [8].

The original Nam models [8] are given for cancer cells as

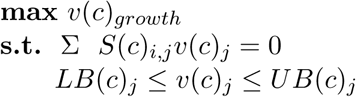

and for normal cells as

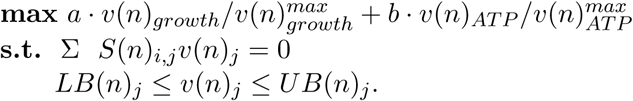

In the above linear programming formalization, *v*(*c*)_*growth*_ and *v*(*n*)_*growth*_ are the growth rate for cancer and normal cells, respectively. 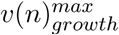 and 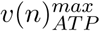 are the upper bounds for the growth rate and ATP production for normal cells, respectively, in Nam models. *S*(*c*) and *S*(*n*) are the stoichiometric matrices for cancer and normal cells, respectively. *v*(*c*)_*j*_ and *v*(*n*)_*j*_ are the sizes of the *j*th flux for cancer and normal cells, respectively. *LB*(*n*)_*j*_ and *UB*(*n*)_*j*_ are the lower and upper bounds for *v*(*n*)_*j*_, and *LB*(*c*)_*j*_ and *UB*(*c*)_*j*_ are the lower and upper bounds for *v*(*c*)_*j*_.

### 5.1 Controllability check for independent Nam models

In the original Nam models the lower bound is −0.05 nmol/gDW/h and the upper bound is some positive value for most exchange uptake reactions. Note that the negative values for exchange reactions correspond positive uptake in Nam models. In our computational experiments, an additional constraint was applied where the ratio of an exchange uptake reaction is limited to a small range. If the resulting growth rate of the cancer cell is less than a given criteria and the resulting growth rate of the normal cell is more than or equal to another criterion, then we consider the designated range of the exchange uptake reaction to be *effective*. The criterion for cancer and normal cells in our computational experimentsare 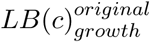 and 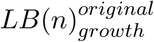, which correspond to their lower bounds of the growth rate in the original Nam models. In the independent models, the computational experiments for cancer and normal cells were conducted independently. The pseudo code for determining whether a range is effective was as follows.

**function** *controllability independent*(*designated reaction, d*)

**for** i=1 **to** d

*lb* = (*i -* 1)*/d*; *ub* = *i/d*;

**max** *v*(*c*)_*growth*_

**s.t.** S *S*(*c*)_*i,j*_*v*(*c*)_*j*_ = 0

*lb · LB*_*j*_ *≤ v*(*c*)_*j*_ *≤ ub · UB*_*j*_ (for *j* = *designated reaction*)

0 *≤ v*(*c*)_*j*_ *≤ UB*(*c*)_*j*_ (for *j* = *growth*)

*LB*(*c*)_*j*_ *≤ v*(*c*)_*j*_ *≤ UB*(*c*)_*j*_ (for *j ≠ designated reaction, growth*)

**max** 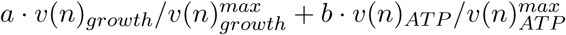

**s.t.** S *S*(*n*)_*i,j*_*v*(*n*)_*j*_ = 0

*lb · LB*_*j*_ *≤ v*(*n*)_*j*_ *≤ ub · UB*_*j*_ (for *j* = *designated reaction*) 0 *≤ v*(*n*)_*j*_ *≤ UB*(*n*)_*j*_ (for *j* = *growth*)

*LB*(*n*)_*j*_ *≤ v*(*n*)_*j*_ *≤ UB*(*n*)_*j*_ (for *j ≠ designated reaction, growth*)

**if** 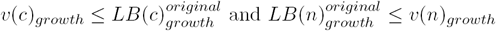

**return** “[*lb · LB*_*j*_, *ub · UB*_*j*_] is effective.”

**else**

**return** “[*lb · LB*_*j*_, *ub · UB*_*j*_] is ineffective.”

### 5.2 Controllability check for integrated Nam models

In the independent model, the models for cancer and normal cells were independently simulated. However, they may affect each other‘s metabolism. To consider such interactions between cancer and normal cells, an integrated model is necessary. In this section, we adapted, to this end, the framework developed in [12] to simulate the interactions between human and microbes.

Fig. 3 (A) and (B) represent toy models of cancer and normal cells, respectively. Circles represent metabolites and arrows represent reactions. 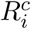 and 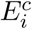 represent internal and exchange reactions in cancer cells, respectively. Similarly, 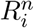 and 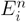 represent internal and exchange reactions in normal cells, respectively. In Fig. 3 (A), seven metabolites, five internal reactions, and three exchange reactions are represented. Similarly, in Fig. 3 (B), seven metabolites, four internal reactions, and three exchange reactions are represented. Fig. 4 shows the stoichiometric matrices for the networks of Fig. 3.

**Figure 3.**
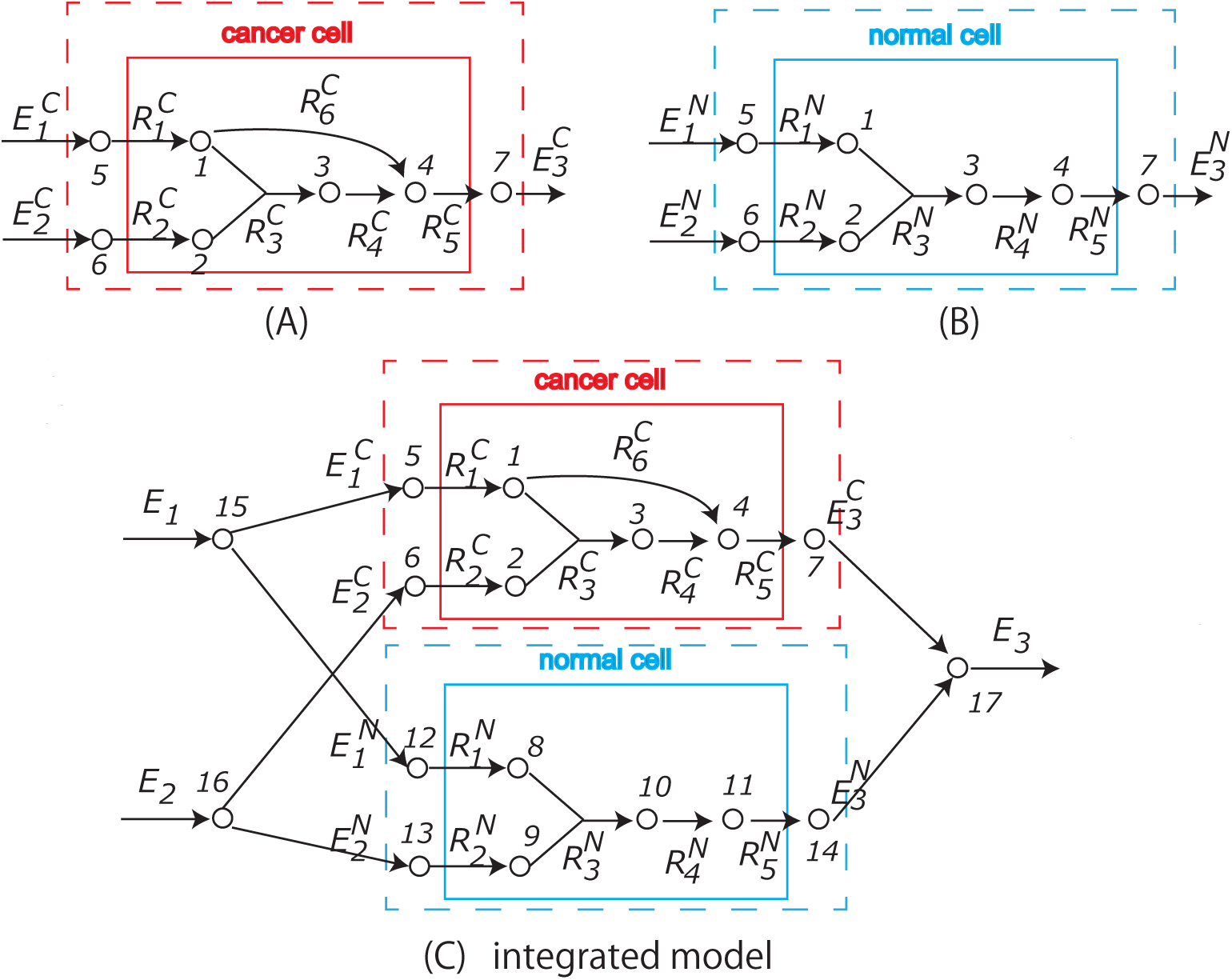
Toy examples of metabolic networks for (A)cancer cells, (B)normal cells, (C)their integrated models.

**Figure 4.**
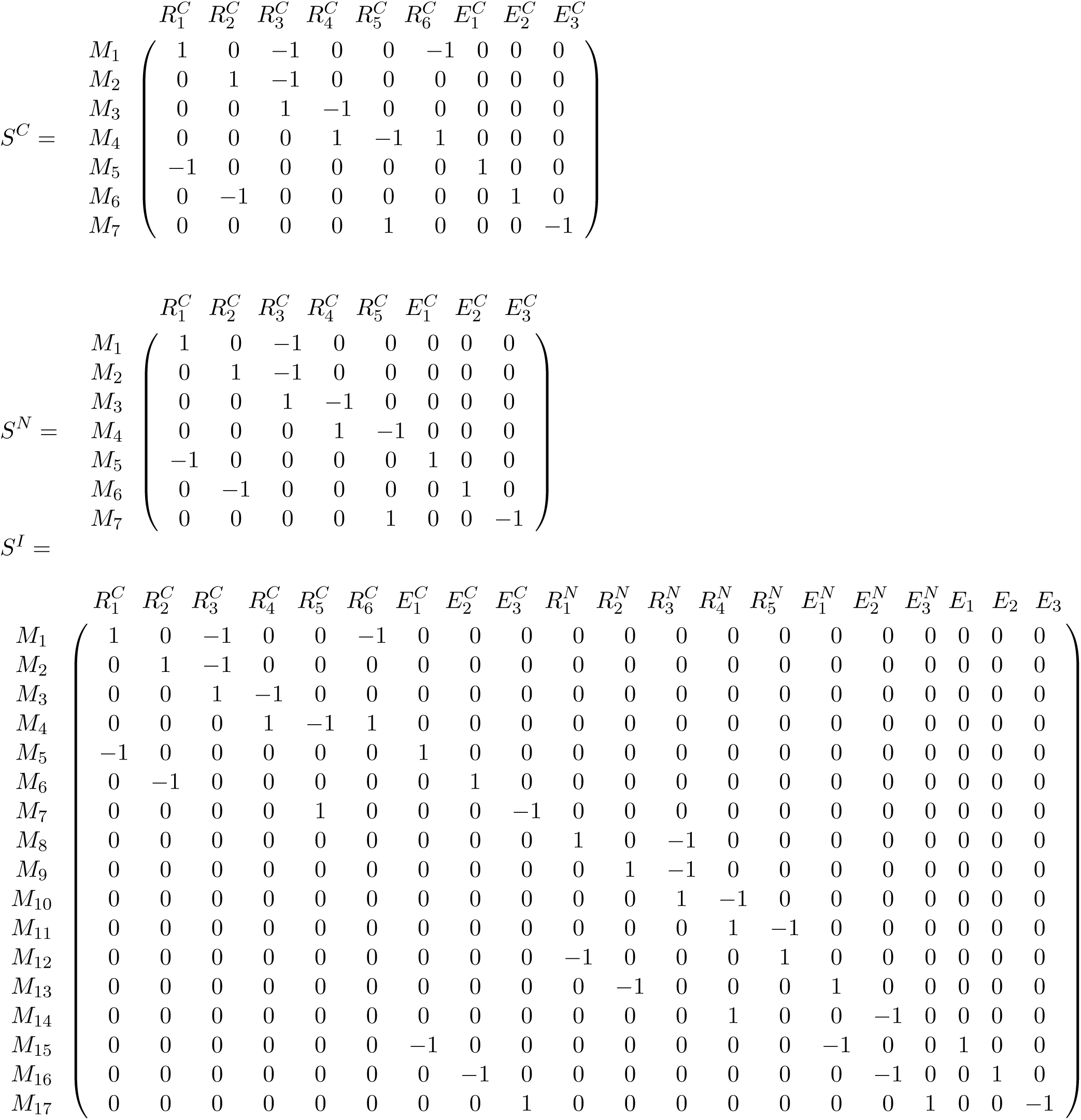
*S*^*C*^, *S*^*N*^, and *S*^*I*^ are stoichiometric matrix for the networks of Fig. 3 (A), (B), and (C), respectively.

The pseudo code for controllability check for the integrated Nam models is as follows.

**function** *controllability integrated*(*designated reaction, d*)

**for** i=1 **to** d

*lb* = (*i -* 1)*/d*; *ub* = *i/d*;

**max** 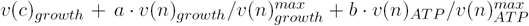

**s.t.** S *S*(*c*)_*i,j*_*v*(*c*)_*j*_ = 0 S *S*(*c*)_*i,j*_*v*(*n*)_*j*_ = 0 *LB*(*c*)_*j*_ *≤ v*(*c*)_*j*_ *≤ UB*(*c*)_*j*_ (for *j ≠*growth) *LB*(*n*)_*j*_ *≤ v*(*c*)_*j*_ *≤ UB*(*n*)_*j*_ (for *j ≠ growth)* 0 *≤ v*(*c*)_*j*_ *≤ UB*(*c*)_*j*_ (for *j* = *growth*) 0 *≤ v*(*n*)_*j*_ *≤ UB*(*c*)_*j*_ (for *j* = *growth*) *lb · LB(ex)*_*j*_ *≤ v*(*ex*)_*j*_ *≤ ub · UB(ex)*_*j*_ (for *j* = *designated reaction*) *LB*(*ex*)_*j*_ *≤ v*(*ex*)_*j*_ *≤ UB*(*ex*)_*j*_ (for *j ≠ designated reaction*)

**if** 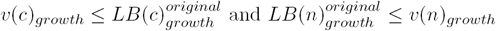

**return** “[*lb · LB*_*j*_, *ub · UB*_*j*_] is effective.”

**else**

**return** “[*lb · LB*_*j*_*, ub · UB*_j_] is ineffective.”

## Data availability

All source codes are included in supplementary information.

## Supplementary information

All source codes are included in supplementary information.

## Acknowledgments

T.T. was supported by grants from JSPS, KAKENHI #16K00391 and #16H02485.

## Author contributions

This work was done only by TT.

## Competing interests

None declared.

## APPENDIX

**Table A1.** Controllability of nine types of cancer by changing the ratio of uptake reactions in integrated Nam models. 1:lung ADC, 2:breast, 3:gastric, 4: kidney, 5: leukemia, 6:liver, 7:lung SCC, 8: ovarian, 9: pancreas.

| ID Ex | EAA | Rxns | 1 | 2 | 3 | 4 | 5 | 6 | 7 | 8 | 9 |
| --- | --- | --- | --- | --- | --- | --- | --- | --- | --- | --- | --- |
| 39 | 1 | L-histidine | 1 | 1 | 1 | 1 | 1 | 1 | 1 | 1 | 1 |
| 40 | 1 | L-Isoleucine | 1 | 1 | 1 | 1 | 1 | 1 | 1 | 1 | 1 |
| 44 | 1 | L-Leucine | 1 | 1 | 1 | 1 | 1 | 1 | 1 | 1 | 1 |
| 46 | 1 | L-Lysine | 1 | 0 | 1 | 0 | 1 | 0 | 0 | 1 | 1 |
| 47 | 1 | L-Methionine | 1 | 1 | 1 | 1 | 1 | 1 | 1 | 1 | 1 |
| 52 | 1 | L-phenylalanine | 1 | 1 | 1 | 1 | 1 | 1 | 1 | 1 | 0 |
| 69 | 1 | L-Threonine | 1 | 1 | 1 | 1 | 1 | 1 | 1 | 1 | 1 |
| 72 | 1 | L-Tryptophan | 1 | 1 | 1 | 1 | 1 | 1 | 1 | 1 | 1 |
| 74 | 1 | L-Valine | 1 | 1 | 1 | 1 | 1 | 1 | 1 | 1 | 1 |
| 18 | conditional | L-Arginine | 1 | 1 | 1 | 1 | 1 | 1 | 1 | 1 | 1 |
| 9 | 0 | 5-Methyltetrahydrofolate | 0 | 0 | 1 | 0 | 1 | 0 | 0 | 1 | 0 |
| 16 | 0 | L-alanine | 0 | 0 | 0 | 0 | 0 | 0 | 0 | 1 | 0 |
| 20 | 0 | L-asparagine | 1 | 0 | 0 | 0 | 1 | 1 | 0 | 0 | 0 |
| 30 | 0 | L-cysteine | 0 | 0 | 1 | 0 | 1 | 0 | 0 | 0 | 0 |
| 32 | 0 | fe2(u) | 0 | 1 | 0 | 0 | 0 | 1 | 0 | 1 | 0 |
| 34 | 0 | D-Glucose | 1 | 0 | 0 | 0 | 1 | 0 | 0 | 0 | 0 |
| 41 | 0 | myo-Inosito | 1 | 0 | 1 | 0 | 1 | 1 | 0 | 1 | 1 |
| 51 | 0 | oxygen | 0 | 0 | 0 | 0 | 0 | 1 | 0 | 1 | 0 |
| 53 | 0 | Phosphate | 0 | 0 | 0 | 0 | 0 | 1 | 0 | 0 | 1 |
| 64 | 0 | L-serine | 0 | 0 | 0 | 0 | 1 | 1 | 0 | 0 | 0 |
| 73 | 0 | L-Tyrosine | 1 | 0 | 0 | 0 | 1 | 0 | 0 | 1 | 0 |

**Table A2.**
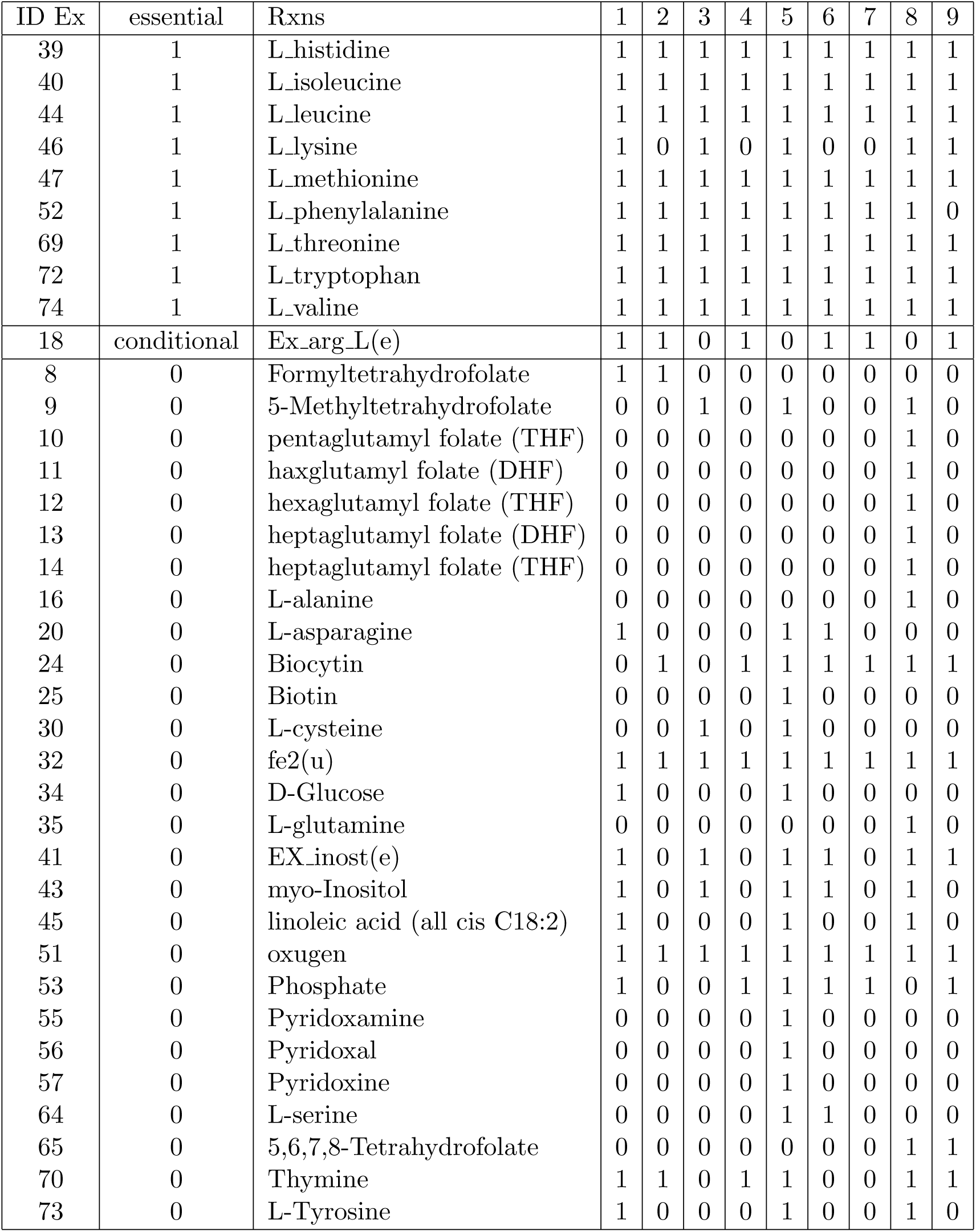
Controllability of nine types of cancer by changing the ratio of uptake reactions for independent Nam models. 1:lung ADC, 2:breast, 3:gastric, 4: kidney, 5: leukemia, 6:liver, 7:lung SCC, 8: ovarian, 9: pancreas

